# Antibiotic-induced decreases in the levels of microbial-derived short-chain fatty acids promote gastrointestinal colonization of *Candida albicans*

**DOI:** 10.1101/428474

**Authors:** Jack Guinan, Shaohua Wang, Hariom Yadav, Shankar Thangamani

## Abstract

*Candida albicans* is the fourth most common cause of systemic nosocomial infections, posing a significant risk in immunocompromised individuals. As the majority of systemic *C. albicans* infections stem from endogenous gastrointestinal (GI) colonization, understanding the mechanisms associated with GI colonization is essential in the development of novel methods to prevent *C. albicans*-related mortality. In this study, we investigated the role of microbial-derived short-chain fatty acids (SCFAs) including acetate, butyrate, and propionate on growth, morphogenesis, and GI colonization of *C. albicans*. Our results indicate that cefoperazone-treated mice susceptible to *C. albicans* infection had significantly decreased levels of SCFAs in the cecal contents that correlate with a higher fungal load in the feces. Further, using *in vivo* concentration of SCFAs, we demonstrated that SCFAs inhibit the growth, germ tube, hyphae and biofilm development of *C. albicans in vitro*. Collectively, results from this study demonstrate that antibiotic-induced decreases in the levels of SCFAs in the cecum enhances the growth and GI colonization of *C. albicans*.

## INTRODUCTION

*C. albicans*, often present in the healthy gastrointestinal (GI) tract, is harmless to the immunocompetent human host with its resident microbiota^1,2^. Though compelling evidence suggests that disturbances in immune regulation contribute to invasive *C. albicans* infections, antibiotic-induced gut dysbiosis remains a major risk factor for increased *C. albicans* colonization and dissemination in immunocompromised patients and individuals with antibiotic-associated diarrhea (AAD)^3–16^. Administration of broad-spectrum antibiotics increases the risk of *C. albicans* colonization in the gut and the source of systemic infections is often found to be the GI tract ^13,16–20^. In addition, more than 60% of individuals with AAD test positive for *C. albicans* and patients treated with antibiotics for *Clostridium difficile* often develop an episode of candidemia^9,10,14,15,21^. Taken together, these studies demonstrate that antibiotic-induced gut dysbiosis in immunocompromised individuals and AAD patients leads to increased colonization of *C. albicans* and this increased intestinal colonization predisposes high-risk patients to systemic candidiasis ^22,23^. Therefore, understanding the factors involved in antibiotic-induced gut dysbiosis and subsequent GI colonization of *C. albicans* is critical to treat and prevent *C. albicans* pathogenesis.

Antibiotic treatment in mice and humans alters the composition of gut microbiota, ultimately leading to changes in the levels of microbial-derived gut metabolites, mainly bile acids and short-chain fatty acids (SCFAs)^24–28^. Alterations in the normal levels of microbial-derived bile acids and SCFAs have been implicated in the growth, colonization, and pathogenesis of enteric pathogens including *C. difficile ^24,25,28^*. Moreover, we have recently demonstrated that microbial-derived bile acids play an important role in the GI colonization of *C. albicans* ^29 30^. However, the role of SCFAs including acetate, propionate, and butyrate–three major fatty acids produced by gut microbiota^31–36^–in the GI colonization of *C. albicans* is poorly understood. Given the abundance of SCFAs in the intestine, a natural habitat and invasion site for *C. albicans*, understanding the role of SCFAs on fungal growth, morphogenesis and colonization will have important implications in *C. albicans* infections. Therefore, in this study, we aim to understand the role of microbial-derived SCFAs in the GI colonization of *C. albicans*.

To investigate if antibiotic treatment alters the levels of microbial-derived SCFAs and GI colonization of *C. albicans*, we treated mice with cefoperazone and determined the levels of SCFAs and the *C. albicans* load in the cecal and fecal contents, respectively. Furthermore, the role of SCFAs including acetic, butyric, and propionic acid on *C. albicans* growth and morphogenesis were investigated *in vitro*. Our results indicate that SCFAs inhibit the growth and morphogenesis of *C. albicans* and may potentially regulate the GI colonization of this fungal pathogen.

## RESULTS

### Antibiotic-treated, *C. albicans-susceptible* mice have significantly reduced levels of SCFAs in the cecum

To determine the impact of antibiotic treatment on cecal SCFA levels and *C. albicans* load, groups of mice were treated with sterile water with or without cefoperazone for 7 days. After 7 days of antibiotic treatment, mice were euthanized for cecal SCFA analysis. Another set of control or antibiotic-treated mice were infected with *C. albicans* and their fecal CFU load was determined after 5 days of infection.

Results indicate that cefoperazone-treated mice had a significantly higher *C. albicans* load in the feces after 5 days of infection. Cefoperazone-treated mice had an almost 3 log_10_ increase in fungal load in the feces compared to control groups (Fig. 1a). Next we determined the SCFA levels in cefoperazone-treated *C. albicans* susceptible group and non-treated control group that are resistant to *C. albicans* infection. Interestingly, SCFA levels in the cecum of cefoperazone-treated mice were significantly decreased compared to control groups (Fig. 1b). The average concentration of acetic acid, butyric acid and propionic acid in the cecal content of control groups were 36.87 ± 7.11 μmol/g, 7.52 ± 0.92 μmol/g and 8.18 ± 0.77 μmol/g respectively. However, the SCFA levels in the cefoperazone-treated mice that are highly susceptible to *C. albicans* were acetic acid (16.13 ± 2.39 μmol/g), butyric acid (1.77 ± 0.79 μmol/g) and propionic acid (1.95 ± 0.63 μmol/g) (Fig. 1b). Taken together, these results suggest that cefoperazone-treated mice susceptible to *C. albicans* GI colonization had significantly decreased levels of SCFAs in the cecal contents.

**Fig. 1.**
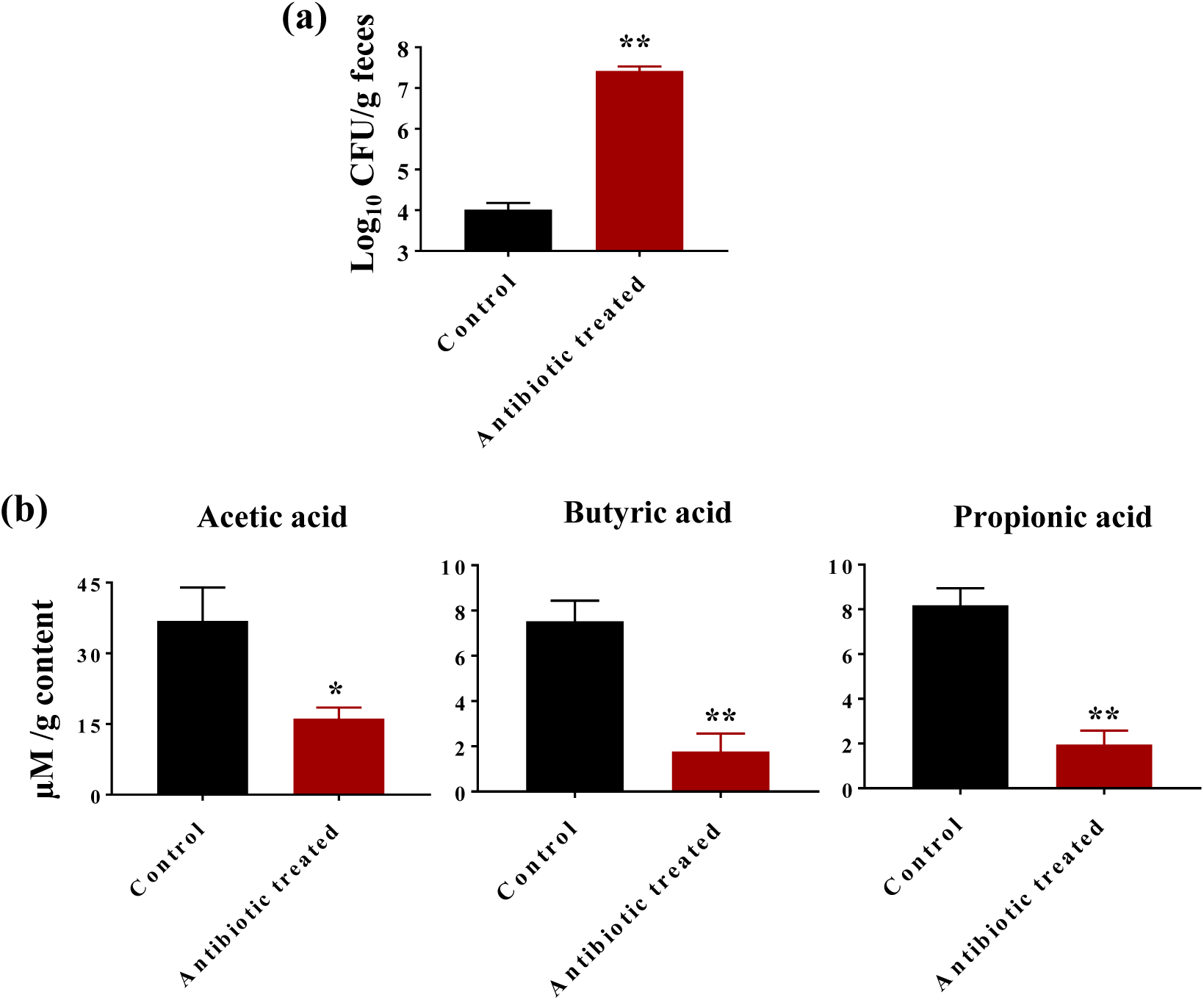
Cefoperazone-treated mice susceptible to *C. albicans* have decreased levels of SCFAs in the cecum. *C. albicans* SC5314 load in fecal contents after 5 days of infection in mice receiving sterile water with or without cefoperazone. Fungal load (Log_10_ CFU/g feces) determined by CFU count (a). SCFA levels (μmol/g) in the cecal contents from mice receiving sterile water with or without cefoperazone (b). Data is represented as means ± SEM with n = 5-6 mice in each treatment group. Statistical significance was evaluated using student’s t-test and *P* values (* ≤ 0.05, ** ≤ 0.01) were considered as significant.

### SCFAs inhibit the growth of *C. albicans in vitro*

To investigate if *in vivo* levels of SCFAs in the cecal contents have any potential role in GI colonization of *C. albicans*, we examined the effect of SCFAs on *C. albicans* growth *in vitro*. We used pH-adjusted RMPI media to determine if changes in pH as a result of SCFA treatment have any inhibitory effect on *C. albicans* growth. RPMI media was titrated with HCl to match the pH values of SCFA treatments (Table 1). Results indicate that *C. albicans* (ATCC 10231 and SC 5314) strains grown in a pH ranging from 7.00 to 3.65±0.05 experienced strain-dependent changes in growth in different pH-adjusted RPMI media (Fig. 2a and 2b). *C. albicans* SC 5314 exhibited a 30% increase in growth at pH 3.65±0.05, 4.12±0.07 and 4.49±0.04, and a 12% increase at pH 5.38±0.05 compared to fungal cells grown in pH 7.00 RPMI control after 24 hours (Fig. 2a). On the other hand, *C. albicans* ATCC 10231 strain did not show a considerable change in growth at pH values ranging from 7.00 to 4.12±0.07. However, it exhibited a 13% decrease in growth at a pH value of 3.65±0.05 compared to pH 7.00 RPMI control (Fig. 2b). These results indicate that alteration in pH does not considerably inhibit the growth of *C. albicans*.

**Fig. 2.**
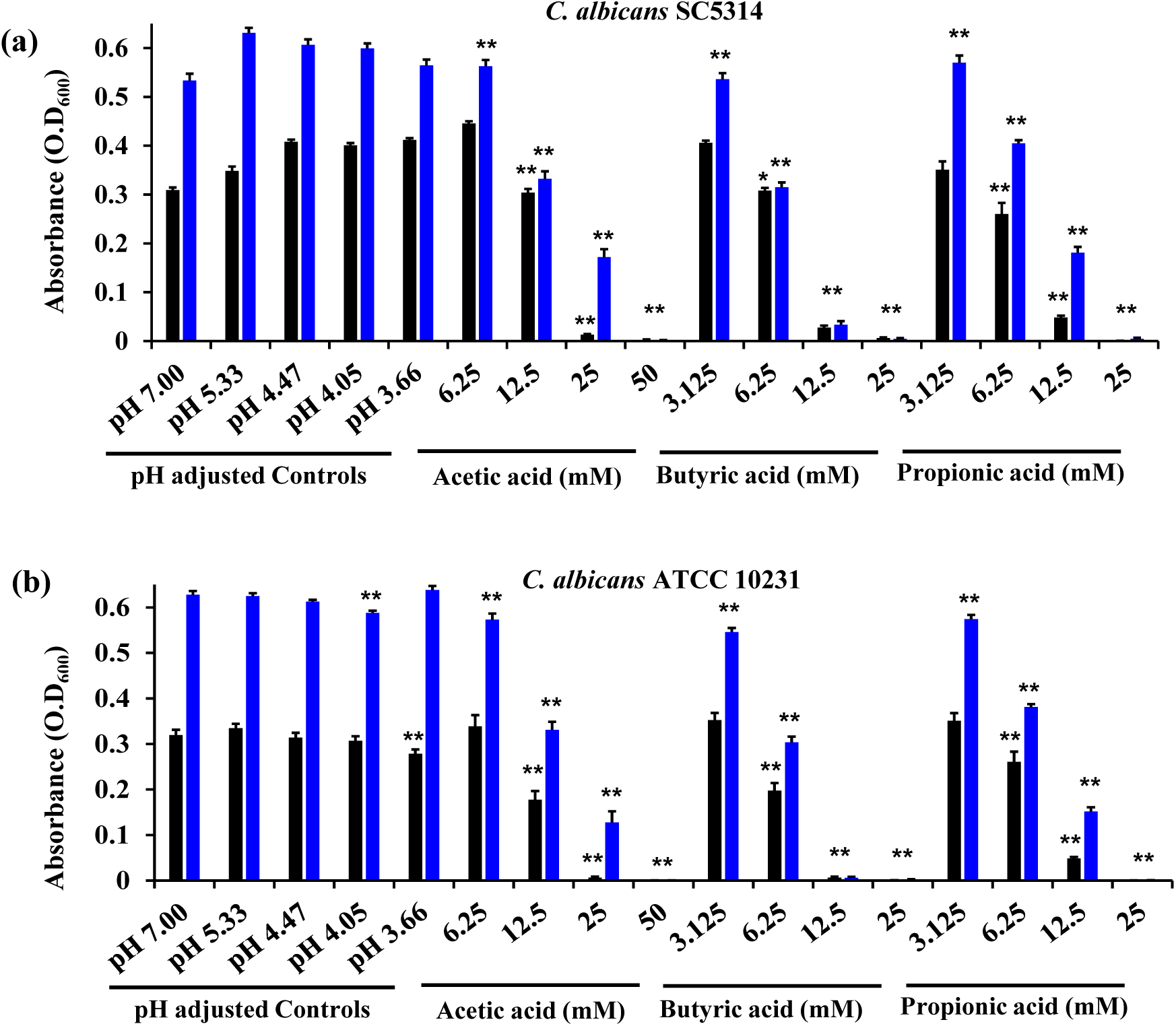
SCFAs inhibit *C. albicans* growth *in vitro*. Growth of *C. albicans* strains SC 5314 (a) and ATCC 10231 (b) in the presence of SCFAs or in pH-adjusted RPMI media determined by spectrophotometer analysis at an optical density of 600 nm after 24 and 48 hours of incubation. Experiment was repeated three times and the three combined replicates were shown here with total n = 9 for each group. Data is represented as means ± SEM. Statistical significance was evaluated using student’s t-test and *P* values (* ≤ 0.05, ** ≤ 0.01) were considered as significant. Significance is shown only for data points that exhibited significant decreases in growth compared to respective controls. Significance for pH-adjusted RPMI values was assessed using pH 7.00 as the comparative data set; significance for SCFA-treated conditions was assessed using respective pH controls for each SCFA condition in statistical analyses. Significance (**) for acetic acid (50 mM), butyric acid (12.5 mM and 25 mM), and propionic acid (25 mM) indicates p ≤ 0.01 at both 24 and 48 hours in both strains.

**Table 1.** Experimental pH of RPMI supplemented with varying concentrations of acetic acid and respective pH of HCl-adjusted pH controls. Values taken as mean ± SEM.

| RPMI (for growth and biofilm Assays) |  |  | RPMI + 30% FBS (for hyphal and germ tube assays) |  |  |
| --- | --- | --- | --- | --- | --- |
| [Acetic Acid] (mM) | Experimental pH | HCl-Adjusted Controls pH | [Acetic Acid] (mM) | Experimental pH | HCl-Adjusted Controls pH |
| 50 | 3.66±0.03 | 3.65±0.05 | 50 | - | - |
| 25 | 4.05±0.03 | 4.12±0.07 | 25 | 4.68±0.02 | 4.69±0.01 |
| 12.5 | 4.47±0.03 | 4.49±0.04 | 12.5 | 5.46±0.05 | 5.52±0.03 |
| 6.25 | 5.33±0.06 | 5.38±0.05 | 6.25 | - | - |
| 3.125 | 6.36±0.03 | 5.85±0.24 | 3.125 | - | - |
| 0 | 7.00 | 7.00 | 0 | >7.00 | 7.01±0.01 |

Next, we determined the effect of SCFAs on *C. albicans* growth. In RPMI media supplemented with average *in vivo* concentrations of SCFAs, *C. albicans* exhibited a significant decrease in growth compared to its respective pH-adjusted control groups (Fig. 2a and 2b). After 24 hours of treatment with acetic acid, *C. albicans* (SC 5314 and ATCC 10231) exhibited a 25–40% decrease at 12.5 mM, a 95% decrease at 25 mM, and no growth at 50 mM (Fig. 2a and 2b). Butyric acid exhibited potent inhibitory activity even at lower concentrations. Butyric acid (6.25 mM) reduced growth by 11–41%, followed by reduction of growth by 93–98% and 98–100% at 12.5 mM and 25 mM treatments, respectively (Fig. 2a and 2b). A similar trend was seen with propionic acid. At a concentration of 6.25 mM, propionic acid exhibited a 22% reduction in growth *C. albicans* (ATCC 10231). *C. albicans* (SC 5314 and ATCC 10231) further exhibited a 57–84% and 98–100% decrease in growth in the presence of 12.5 mM and 25 mM of propionic acid, respectively (Fig. 2a and 2b). These trends continued into 48 hours, with strain-dependent significant inhibition of growth at varying concentrations of acetic acid (12.5 – 50 mM), butyric acid (6.25 – 25 mM), and propionic acid (6.25 – 25 mM) (Fig. 2a and 2b). Overall, our results demonstrate that SCFAs inhibit the growth of *C. albicans* strains in a concentration-dependent manner and that growth inhibition is not due to changes in pH.

### SCFAs inhibit *C. albicans* germ tube formation

The impact of SCFA treatment on *C. albicans* germ tube formation was determined using microscopy analysis. pH-adjusted RPMI controls were used to determine if alterations in pH have any effect on germ tube formation. Microscopic imaging revealed that *C. albicans* considerably reduced the germ tube formation in pH controls for 12.5 mM (pH 5.52±0.03) and 25 mM (pH 4.69±0.01) compared to the RPMI control (pH 7.01±0.01) (Fig. 3a). Quantification of the *C. albicans* cells that formed germ tubes after 2 hours of incubation revealed a 30% decrease in germ tube formation in pH control 12.5 mM (pH 5.52±0.03) and an almost 50% decrease in pH control 25 mM (pH 4.69±0.01) compared to the RPMI control (pH 7.01±0.01) (Fig. 3b). These results indicate that acidic pH significantly inhibited the germ tube formation in *C. albicans*.

**Fig. 3.**
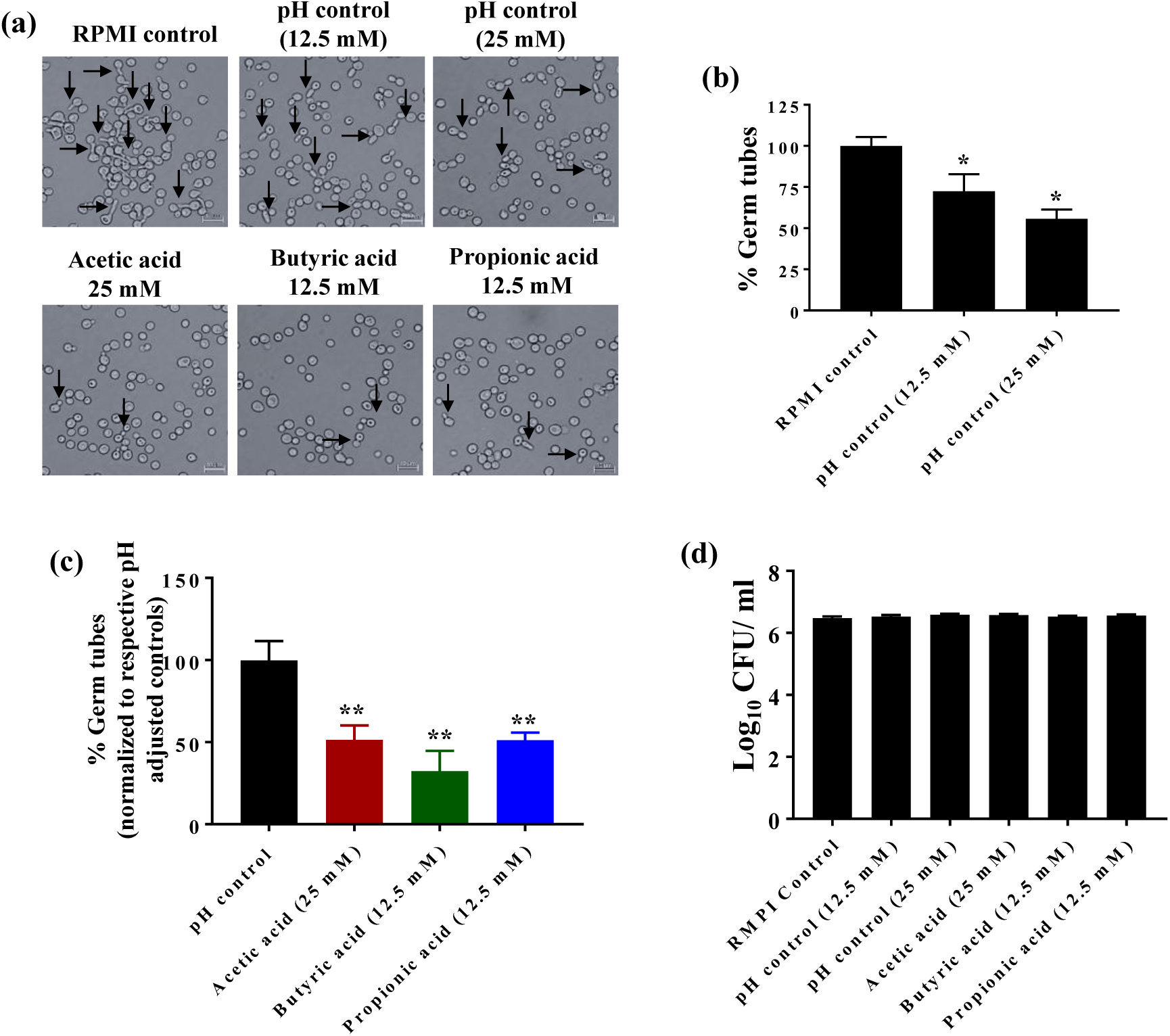
SCFAs inhibit germ tube formation. Germ tube formation in *C. albicans* ATCC 10231 strain in the presence of SCFAs or pH-adjusted media supplemented with 30% FBS. Representative images of germ tubes formed after 2 hours in control and treatment groups determined by microscopic analysis at 40X magnification (a). Quantification of the percent *C. albicans* cells with germ tubes in pH-adjusted controls; pH-adjusted controls (12.5 mM and 25 mM) were normalized to the RPMI control (pH 7.00) (b). Quantification of the percent *C. albicans* cells with germ tubes in SCFA treatment groups (c). SCFA treatments were normalized to their respective pH controls. The germ tube experiment was repeated three times and two 40X images were taken from each replicate for each treatment group, with a minimum n = 1000 cells for each group. *C. albicans* (CFU/mL) viability determined after 2 hours of incubation in germ tube-inducing conditions (d). The experiment was repeated three times with n = 12 for each treatment group. Combined replicates for both experiments are shown here. Data is represented as means ± SEM. Statistical significance was evaluated using student’s t-test and *P* values (* ≤ 0.05, ** ≤ 0.01) were considered as significant.

Although acidic pH itself was shown to be a factor in inhibiting germ tube formation, we determined the effect of SCFAs on *C. albicans* germ tube formation. Interestingly, SCFAs were more potent in inhibiting the germ tube formation compared to their respective pH-adjusted controls. Microscopic imaging of *C. albicans* in the presence of SCFAs revealed a considerable decrease in germ tube formation compared to their respective pH-adjusted controls (Fig. 3a). In addition, quantification of the percentage of germ tubes formed revealed that acetic acid (25 mM) reduced germ tube formation by 50% compared to its pH control (pH 4.69±0.01) (Fig. 3c). Butyric acid (12.5 mM) and propionic acid (12.5 mM) also significantly inhibited germ tube formation compared to their pH control (pH 5.52±0.03), reducing germ tube formation by nearly 70% and 50%, respectively (Fig. 3c).

To determine if germ tube inhibition by SCFAs was not due to fungal cell death, *C. albicans* cells incubated in the germ tube conditions were determined for cell viability. Results indicated that no significant decrease in fungal cells was noticed after 2 hours of incubation in SCFA-treated or pH-adjusted control groups (Fig. 3d). These results indicate that SCFAs inhibit *C. albicans* germ tube formation partly by inducing acidic conditions and through other unknown mechanisms.

### SCFAs inhibit *C. albicans* hyphae formation

The effect of SCFAs on *C. albicans* hyphae formation was evaluated using crystal violet and microscopic analyses. We used pH-adjusted RMPI media to determine if changes in pH have any effect on *C. albicans* hyphae formation. Results indicate that *C. albicans* grown in RPMI media at pH 7.01±0.01 (RPMI control) showed massive hyphae formations (Fig. 4a). However, a considerable decrease in hyphae formation was noticed in the pH-adjusted controls for 12.5 mM (pH 5.52±0.03) and 25 mM (pH 4.69±0.01) treatments (Fig. 4a). Further, a crystal violet assay indicated that the pH-adjusted control 25 mM (pH 4.69±0.01) inhibited 90% of *C. albicans* hyphae attachment, followed by 75% inhibition in the pH-adjusted control 12.5 mM (pH 5.52±0.03) (Fig. 4b). Taken together, the crystal violet assay complements the findings of microscopic observations, indicating that acidic pH not only decreases hyphae formation but also significantly inhibits *C. albicans* hyphae attachment to polystyrene plates.

**Fig. 4.**
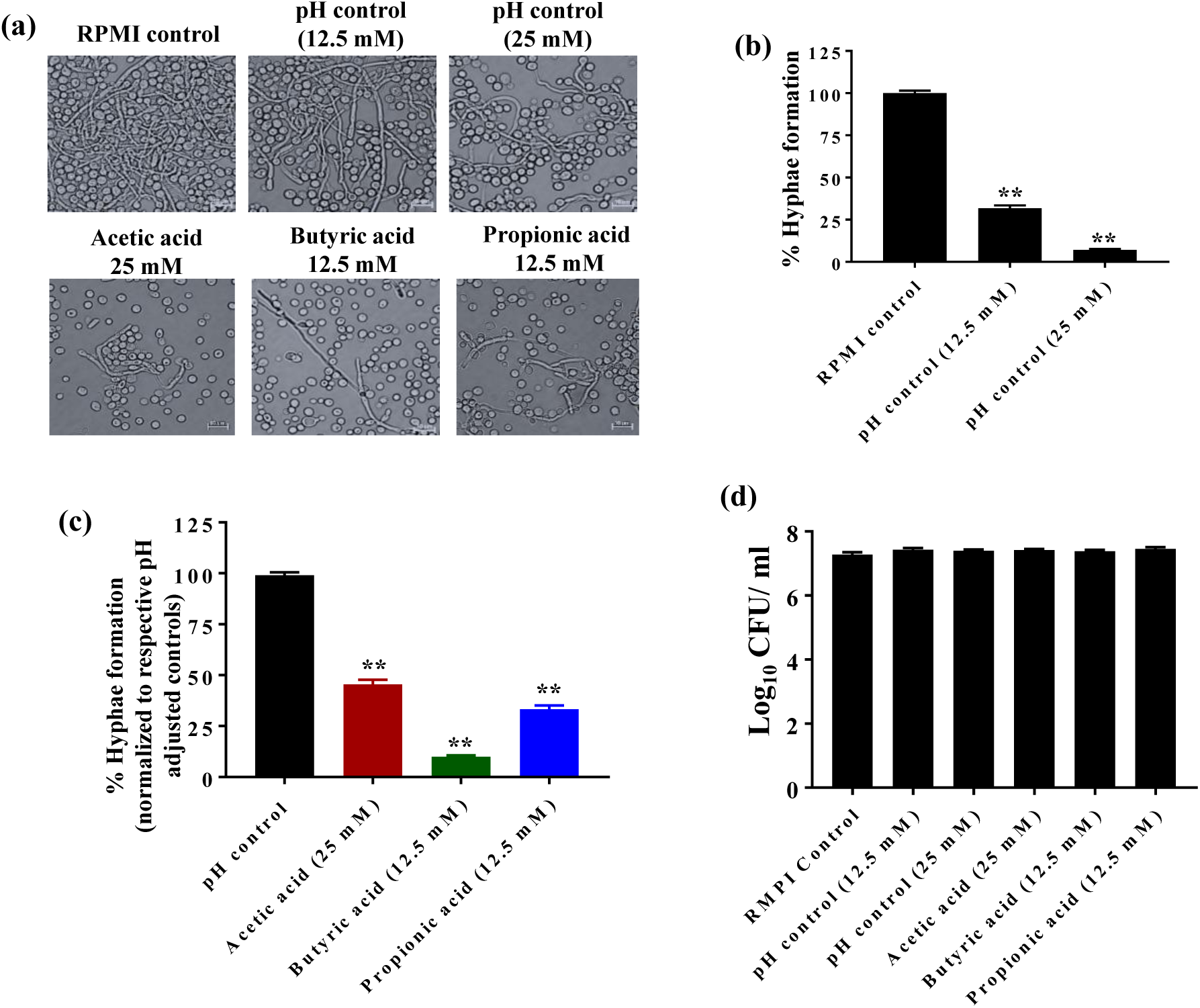
SCFAs inhibit *C. albicans* hyphae formation *in vitro*. *C. albicans* ATCC 10231 was grown in the presence of SCFAs or in pH-adjusted RPMI media supplemented with 30% FBS and examined using bright field microscopy at 40X (a). Quantification of *C. albicans* hyphae attachment to polystyrene plates in pH-adjusted controls; pH-adjusted controls (12.5 mM and 25 mM) were normalized to the RPMI control (pH 7.00) (b). Quantification of *C. albicans* hyphae attachment to polystyrene plates in SCFA-treatment groups; SCFA treatment groups were normalized to their respective pH controls (c). *C. albicans* (CFU/mL) viability determined after 12 hours of incubation in hyphae-inducing conditions (d). The experiment was repeated three times with n = 24 for the hyphae formation and n = 12 for the CFU viability determination in each treatment group. Data is represented as means ± SEM. Statistical significance was evaluated using student’s t-test and *P* values (* ≤ 0.05, ** ≤ 0.01) were considered as significant.

Next, we investigated the effect of SCFAs on *C. albicans* hyphae formation and attachment. Our results indicate that SCFAs including acetic acid (25 mM), butyric acid (12.5 mM) and propionic acid (12.5 mM) considerably decreased *C. albicans* hyphae formation compared to their respective pH-adjusted RPMI controls (Fig. 4a). Further, all three SCFAs significantly inhibited *C. albicans* hyphae attachment to the polystyrene plates (Fig. 4c). Butyric acid (12.5 mM) inhibited 90% of *C. albicans* hyphae attachment, followed by propionic acid (12.5 mM) and acetic acid (25 mM) by 70 and 50%, respectively compared to their pH-adjusted controls (Fig. 4c).

To rule out if *C. albicans* hyphae inhibition as a result of SCFA treatment was not due to fungal cell death, *C. albicans* grown under hyphae-inducing conditions in the presence or absence of SCFAs were plated onto YPD agar plates to determine the CFU count. Results from this experiment suggest that viability of fungal cells was not significantly affected in all treatment conditions compared to RPMI control (pH 7.01±0.01), indicating that decreased hyphae formation and attachment was not due to decrease in cell viability (Fig. 4d). Taken together, our results indicate that SCFAs may regulate *C. albicans* hyphae formation and attachment partly by altering pH levels in addition to other mechanisms.

### SCFAs reduce the metabolic activity of fungal cells in *C. albicans* biofilm

The effect of SCFAs on the metabolic activity of *C. albicans* ATCC 10231 in the biofilm was evaluated using an MTS reduction assay. In order to determine if the change in pH has any effect on the metabolic activity of fungal cells in *C. albicans* biofilm, we used pH-adjusted RMPI media to determine the effect of pH on *C. albicans* metabolic activity in the biofilm. Results indicate that an acidic pH significantly decreases the metabolic activity of fungal cells in the biofilm (Fig. 5a). The metabolic activity of fungal cells in *C. albicans* biofilm in the pH-adjusted controls 12.5 mM (pH 5.52±0.03) and 25 mM (pH 4.69±0.01) were decreased by 25% compared to the RPMI control (pH 7.00) (Fig. 5a).

**Fig. 5.**
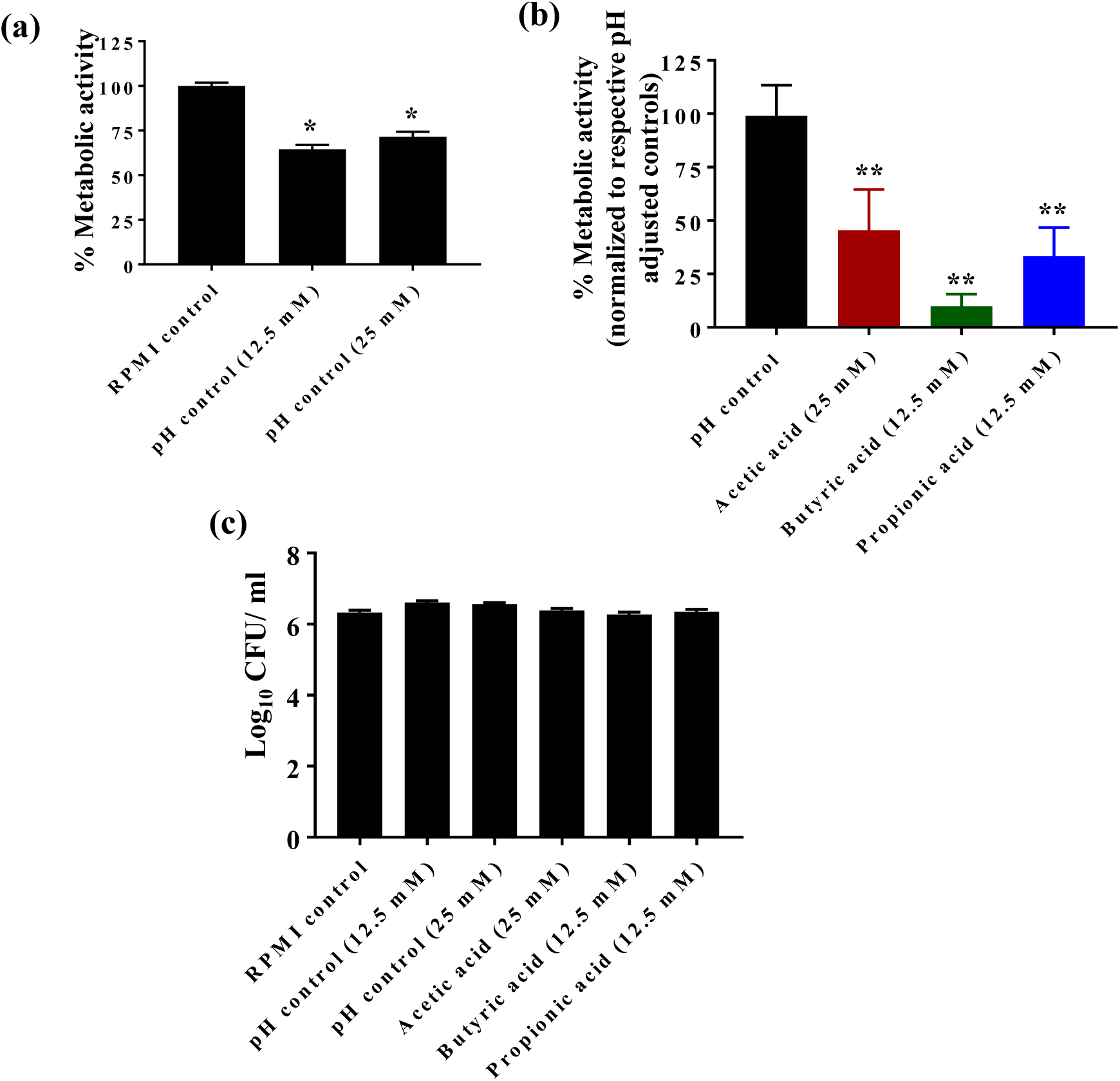
SCFAs reduce the metabolic activity of fungal cells in the *C. albicans* biofilm. *C. albicans* ATCC 10231 was grown in the presence of SCFAs or in pH-adjusted RPMI and the metabolic activity of the fungal cells in the biofilm was assessed using MTS assay. Percent metabolic activity of fungal cells in the biofilm formed in pH-adjusted controls was determined; pH adjusted controls (12.5 mM and 25 mM) were normalized to the RPMI control (pH 7.00) (a). Percent metabolic activity of fungal cells in the biofilm formed in SCFA-treatment groups; SCFA treatments groups were normalized to their respective pH controls (b). *C. albicans* (CFU/mL) viability determined after 48 hours of incubation in biofilm-inducing conditions (c). All experiments were repeated three times, with n = 18 to determine the metabolic activity in the biofilm and n = 12 for the CFU viability analysis in each treatment group. Data is represented as means ± SEM. Statistical significance was evaluated using student’s t-test and *P* values (* ≤ 0.05, ** ≤ 0.01) were considered as significant.

Next, we assessed the effect of SCFAs on fungal cell metabolic activity in *C. albicans* biofilm. Results indicate that SCFAs including acetic acid (25 mM), butyric acid (12.5 mM) and propionic acid (12.5 mM) significantly decreased the metabolic activity of fungal cells in *C. albicans* biofilm compared to their respective pH-adjusted RPMI controls (Fig. 5b). Butyric acid at 12.5 mM decreased the metabolic activity by 90%, followed by propionic acid (12.5 mM) and acetic acid (25 mM) by 70% and 60%, respectively (Fig. 5b). Further, in order to determine if the effect of SCFAs or pH-adjusted RPMI groups on *C. albicans* metabolic activity was not due to fungal cell death, *C. albicans* grown under biofilm-inducing conditions were plated onto YPD agar plates to determine the CFU count. Results indicate that the viability of fungal cells was not significantly affected in any treatment condition compared to RPMI control (pH 7.00), suggesting that decreased metabolic activity of fungal cells in *C. albicans* biofilm was not due to cell viability (Fig. 5c). Collectively, results from this experiment indicate that SCFAs regulate the metabolic activity of *C. albicans* in the biofilm partly by altering pH and through other mechanisms.

## DISCUSSION

The gut microbiota plays a major role in the colonization resistance to enteric bacterial and fungal pathogens including *C. albicans*^37–45^. While mechanisms of colonization resistance to enteric pathogens by commensal bacteria are speculated to include immune responses, competition for nutrients, pH modulation, and synthesis of antimicrobial and antifungal compounds ^29,37–39,44–51^, the mechanisms associated with colonization resistance to *C. albicans* remain poorly understood. The development of effective preventative and therapeutic treatments thus necessitates a deeper understanding of the innate mechanisms of colonization resistance to *C. albicans*.

Commensal bacteria produce a variety of bioactive molecules; however, SCFAs have emerged as key regulators of gut homeostasis for colonization resistance against enteric pathogens^25,28,33,44^. Several preliminary studies have highlighted the antifungal potential of select SCFAs, including the inhibition of *C. albicans* germination by butyric acid^52,53^, a dose-dependent fungicidal effect with acetic, butyric, and propionic acid treatment^2,54^, an increase in programmed cell death with acetic acid^55^, an increase in mitochondrial-mediated apoptosis with propionic acid^56^, and an upregulation of transcriptional stress responses with acetic, butyric, and propionic acids^54,57^. However, using *in vivo* concentrations, the role of SCFAs on *C. albicans* growth, morphogenesis and GI colonization is poorly understood. Importantly, though the inhibitory effects of SCFAs on enteric pathogens has been speculated to occur due to alterations in pH levels, this hypothesis has not been investigated in detail, particularly in *C. albicans*^58,59^

SCFAs including acetate, butyrate and propionate are produced as a result of bacterial fermentation in the cecum^32,33,36,60,61^. SCFAs are an abundant microbiota metabolites in the GI tract luminal microenvironment, where *C. albicans* colonization takes place^19,20,62^. The molar ratio of acetate, butyrate, and propionate is typically reported as 60:20:20 in the human intestinal tract^63,64^. While some studies report slight differences in this ratio in mice, the trend remains that acetate is the predominate SCFA present in the cecum, followed by butyrate and propionate in varying amounts ^65–67^ In mice (cecum) and humans (feces), the concentration of acetate ranges from 30.09 ± 2.09 − 40.66 ± 0.122 μmol/g to 69.1 ± 5.0 − 73.7 ± 21.5 μmol/g^63,65–67^. Butyrate concentrations in mice and humans remain relatively consistent, ranging from 18.52 ± 4.92 − 35.9 ± 10.2 μmol/g, although concentrations as low as 2.59 ± 0.31 μmol/g have been reported^63,65–67^. Propionate has been reported to range from 7.43 ± 0.16 − 25.3 ± 3.7 μmol/g^63,65–67^. In this study, we also report that cecal SCFA concentrations in control mice are as follows: acetic acid (36.87 ± 7.11 μmol/g), butyric acid (7.52 ± 0.92 μmol/g), and propionic acid (8.18 ± 0.77 μmol/g), which are in agreement with the previous findings^63,65–67^. Since antibiotic-induced gut dysbiosis is an important factor for the GI colonization of *C. albicans*^7,16,68,69^, we examined if antibiotic-induced alterations in the levels of SCFAs play a role in the GI colonization of this fungal pathogen. In this study, we found that cefoperazone-treated mice susceptible to *C. albicans* infection had significantly decreased levels of SCFAs in the cecal contents. The concentration of acetic acid in the antibiotic-treated mice was 16.13 ± 2.39 μmol/g, followed by propionic acid (1.95 ± 0.63μmol/g), and butyric acid (1.77 ± 0.79μmol/g). The concentration of acetic acid was decreased by 50%, followed by butyrate and propionate (75%) after 7 days of cefoperazone treatment, which is in agreement with previous findings indicating that antibiotic treatment significantly decreases the levels of SCFAs in the cecum ^25,70,12^. Taken together, antibiotic-induced decreased in the levels of SCFAs may potentially contribute to the GI colonization of *C. albicans*.

Next, we determined to examine if alterations in the levels of SCFAs as a result of antibiotic treatment actually play a role in the growth and morphogenesis of *C. albicans*, leading to increased GI colonization in cefoperazone-treated mice. Using relevant *in vivo* concentrations of SCFAs found in the cecum of control and cefoperazone-treated mice, we examined the effect of SCFAs on the growth of *C. albicans in vitro*. While the reported cecal concentrations of SCFAs vary considerably based on different factors including detection methods in addition to the age and diet of mice^73,74^, we decided to use the average *in vivo* concentrations of SCFAs for the *in vitro* assays that correlate to published *in vivo* levels found in the cecum of mice and humans^63,65–67^. Our results indicate that average *in vivo* concentration of SCFAs (acetic acid: 25 mM, butyric acid: 12.5 mM, and propionic acid: 12.5 mM)^63,65–67^ found in the control mice exhibit significant inhibitory effects on *C. albicans* growth *in vitro*. However, using average *in vivo* concentrations of SCFAs found in the antibiotic-treated mice (acetic acid: 12.5 mM, butyric acid: 3.25 mM, and propionic acid: 3.25 mM)^25,70–72^ in the *in vitro* assays had only minimal effects on the growth of *C. albicans*. The ability of *C. albicans* to cause infection is associated with its morphological switching from yeast to virulent hyphae, which allows the organism to attach, invade and cause disease^75–81^. Therefore, inhibition of the morphological plasticity of *C. albicans* would substantially reduce its pathogenic potential^76,77,82–84^. A variety of factors in the gut, including n-acetylglucosamine and bacterial peptidoglycan, regulate hyphae formation in *C. albicans*^85–87^. Therefore, using *in vivo* concentration of SCFAs, we investigated the effect of SCFAs on *C. albicans* morphogenesis. Our results indicate that acetic acid, butyric acid, and propionic acid at *in vivo* cecal concentrations found in *C. albicans*-resistant mice significantly inhibited the morphogenesis of *C. albicans in vitro*^63,65–67^ Collectively, our *in vivo* and *in vitro* results along with previous findings demonstrate that antibiotic treatment significantly decreased the levels of SCFAs in the cecum and, in turn may allow *C. albicans* to grow and colonize in the gut^25,70–72^.

Next, we investigated if inhibitory effect of SCFAs on *C. albicans* growth and morphogenesis is due to changes in pH levels. SCFAs inducing an acidic environment due to their dissociative properties may be a factor for their inhibitory effect on *C. albicans* growth and morphogenesis. In order to determine if changes in pH have any effect on *C. albicans* growth and morphogenesis, we used HCl to adjust the pH of RPMI media corresponding to the pH values as a result of SCFA dissociation. Interestingly, our results demonstrated that RPMI control media adjusted for pH (from 7.00 to 3.65±0.05) does not have a major inhibitory effect on *C. albicans* growth. However, we found that an acidic pH (4.12±0.07 to 5.52±0.03) significantly inhibits *C. albicans* morphogenesis. These results suggest that the SCFA-mediated inhibition of *C. albicans* growth was not likely due to an alteration in pH levels. However, SCFAs may potentially regulate the morphogenesis of *C. albicans* by inducing an acidic environment. Since our results also indicate that SCFAs further inhibited *C. albicans* morphogenesis compared to their respective pH-adjusted RPMI control groups, other unknown mechanisms may be involved in the SCFA-induced inhibition of *C. albicans* morphogenesis. Taken together, these results indicate that the inhibitory effects of SCFAs on *C. albicans* growth and morphogenesis are not limited to pH alterations, but by other mechanisms that have yet to be elucidated. Therefore, future studies to understand the mechanisms behind the SCFA-mediated effects on *C. albicans* is important to understand its pathogenesis and to develop novel therapeutic approaches.

Diverse microbes in the gut possess the ability to produce SCFAs. Among these, Bacteroides, Ruminococcaceae, Lachnospiraceae, Clostridia, Prevotella, Oscillospira and Verrucomicrobia (*Akkermansia muciniphila*), and Faecalibacteria are most commonly associated with the production of SCFAs^33,34,36,61,88–91^. Fan et al. observed that bacteroides including *Blautia producta* and *Bacteroides thetaiotaomicron* directly affect *C. albicans* colonization through SCFAs^2^. Our recent studies found that the probiotics interventions increase the SCFAs production by modulating human and mice gut microbiome^92^. Therefore, future studies including characterization of probiotic or commensal bacteria to enhance the abundance of SCFA levels may form a novel approach to prevent and treat *C. albicans* colonization and subsequent pathogenesis.

Antibiotic treatment significantly alters SCFA levels in the gut; however, the composition and concentration of other critical gut metabolites including bile acids are also affected^24,25,29,30^. Recently, we have shown that bile acids play an important role in controlling *C. albicans* growth and morphogenesis ^29,30^. We previously demonstrated that treatment with *in vivo* concentrations of the primary conjugated bile acid taurocholic acid (TCA) (0.0125%) promotes *C. albicans* growth and morphogenesis, whereas treatment with *in vivo* concentrations of secondary bile acids deoxycholic acid (DCA [0.05%]) and lithocholic acid (LCA [0.01%]) inhibit *C. albicans* growth and morphogenesis *in vitro* ^29,30^. Therefore, understanding the role of various gut metabolites in the GI colonization of *C. albicans* will expand our knowledge on *C. albicans* pathogenesis. In addition, this may form a strong foundation for efforts to use commensal bacteria to modulate gut metabolites to prevent and treat *C. albicans* infections.

## MATERIALS AND METHODS

### Strains and Reagents

*Candida albicans* ATCC 10231 was purchased from ATCC. *Candida albicans* SC5314 was kindly provided by Dr. Andrew Koh from University of Texas Southwestern Medical Center.^2^ Media used in this study included RPMI 1640 (Gibco, MA), MOPS (Sigma, MO), YPD Agar (BD Biosciences, CA), and Fetal Bovine Serum (Atlanta Biologicals, GA). Short-chain fatty acids (acetic acid, butyric acid, and propionic acid) were purchased from Sigma Aldrich (MO). Mice were purchased from The Jackson Laboratory (ME). Cefoperazone was purchased from Sigma Aldrich (MO). Other materials were purchased as indicated: mouse oral gavages (Kent Scientific, MA), vancomycin (Alfa Aesar, MA), gentamicin (Acros Organics, NJ), paraformaldehyde (Alfa Aesar, MA), and glycerol (DOT Scientific, MI).

### Cefoperazone treatment and *C. albicans* infection in mice

Female C57BL/6J mice (5–6 mice per group) were supplemented with sterile water with or without cefoperazone (0.5 mg/mL)^93^. Cefoperazone water was replaced every two days. After 7 days of antibiotic treatment, mice were either sacrificed for SCFA metabolite analysis or infected with *C. albicans* SC5314 via oral gavage at a dose of 4.25 × 10^8^ CFU per mice^2^. After 5 days of infection, fecal samples were collected from individual mice to determine the fungal load. Briefly, fecal pellets were weighed and homogenized in PBS and the supernatant was plated onto YPD agar plates containing 0.1 mg/mL gentamicin and 0.010 mg/mL vancomycin^2^. After 24 hours of incubation, the colonies were counted and the fungal load (CFU/gram) was determined for individual mice. The Institutional Animal Care and Use Committee (IACUC) at Midwestern University approved this study under MWU IACUC Protocol #2894. The MWU animal care policies follow the Public Health Service (PHS) Policy on Humane Care and Use of Laboratory Animals and the policies laid out in the Animal Welfare Act (AWA). Trained animal technicians performed animal husbandry in our IACUC monitored animal care facility in the Foothills Science Building.

### Quantifying SCFAs levels in the cecal content

Equal amount of snap frozen cecal content was weighed, dissolved in HPLC grade water and supernatants were collected after centrifugation (12,000 g, 10 min), while processing on ice. Concentrations of SCFAs (acetate, propionate and butyrate) were determined using a HPLC system (Waters-2695 Alliance HPLC system, Waters Corporation, Milford, MA, USA) equipped with Aminex HPX-87H column (Bio-Rad Laboratories, Hercules, CA) and DAD detector (210 nm), and eluting with H_2_SO_4_ (0.005 N) mobile phase with a flow rate of 0.6 ml/min at 25°C, as described elsewhere^92^.

### Growth assay

The growth of *C. albicans* ATCC 10231 and SC 5314 was measured as previously described,^29^ using pH-adjusted controls titrated with HCl as described below (Table 1).

### pH-adjusted control RPMI media for *in vitro* assays

pH-adjusted controls were used for all *in vitro* experiments involving SCFAs. Briefly, pH was measured using a Fisherbrand Accumet AE150 pH meter (Thermo Fisher, MA). The pH meter was calibrated each time before use using Orion calibration buffers (Thermo Fisher, MA). pH was adjusted for short-chain fatty acid treatments using acetic acid as the reference point since acetic, butyric, and propionic acid share similar experimental pK_a_s (4.76, 4.83, and 4.87, respectively^94^), with acetic acid having the lowest pK_a_ and thus most potent effect on pH via dissociation.^94^ pH was adjusted using HCl for *in vitro assays* as described in Table 1. For growth and biofilm assays, RPMI media was adjusted with HCl to match the pH values of SCFAs. Similarly, for hyphae and germ tube assays, RPMI media supplemented with 30% FBS was adjusted with HCl as described in Table 1.

### Biofilm assay

*C. albicans* ATCC 10231 with an inoculum size of 1.7 × 10^6^ − 3.2 × 10^6^ CFU/mL was used to form the biofilm and the metabolic activity of fungal cells in the biofilm was carried out using MTS assay as previously described^29^. The effect of SCFAs on *C. albicans* cell viability was assessed by incubating fungal cells with the indicated concentration of SCFAs at 37°C for 48 hours in 4-mL tubes. Appropriate pH-adjusted RPMI control media was used as described in Table 1. After 48 hours of incubation, the cell suspension was plated onto YPD agar plates and the CFUs were counted to determine the effect of SCFAs on *C. albicans* cell viability.

### Germ tube and hyphae assays

The effect of SCFAs on *C. albicans* germ tube and hyphae formation was assessed as previously described ^29^. For the germ tube assay, *C. albicans* ATCC 10231 strain with an inoculum size of 3.48 × 10^6^ CFU/mL was incubated with or without SCFAs for 2 hours and the percentage of germ tubes formed were quantified at 20X magnification ^29^. Similarly, *C. albicans* ATCC 10231 strain (3.34 × 10^7^ − 4.60 × 10^8^ CFU/mL) was incubated with or without SCFAs and the *C. albicans* hyphae formation and attachment were determined using bright field microscopy and crystal violet assay as described before ^29^. The crystal violet assay was adopted using a protocol defined by Abe et al.^95,96^ Further, the effect of SCFAs on *C. albicans* cell viability under germ tube and hyphae assay conditions was assessed by incubating fungal cells with the indicated concentration of SCFAs at 37°C for 48 hours in 4-mL tubes. After 48 hours of incubation, cell suspensions were plated onto YPD agar plates and the CFUs were counted to determine the effect of SCFAs on *C. albicans* cell viability. Appropriate pH-adjusted RPMI control media was used as described in Table 1.

### Statistical analyses

The Student t-test was utilized for statistical analyses using GraphPad Prism 6.0 (GraphPad Software, La Jolla, CA) with *p*-values of (* ≤ 0.05, ** ≤ 0.01) being considered significant.

## Consent for publication

Not applicable.

## Availability of data and materials

The datasets used and/or analyzed during the current study are available from the corresponding author on reasonable request.

## Authors Contributions

**JG:** Designed and conducted experiments, compiled and analyzed data; **SW** and **HY:** Carried out SCFAs analysis; **ST:** Conceived the idea, help to design experiments, analyze and interpret data, and overall supervision of the project; **JG, SW, HY and ST:** Reviewed data and wrote the manuscript.

## Acknowledgements

We thank Dr. Andrew Koh from University of Texas Southwestern Medical Center for providing *Candida albicans* SC5314 strain.

